# Structure reveals mechanism of CRISPR RNA-guided nuclease recruitment and anti-CRISPR viral mimicry

**DOI:** 10.1101/453720

**Authors:** MaryClare F. Rollins, Saikat Chowdhury, Joshua Carter, Sarah M. Golden, Heini M. Miettinen, Andrew Santiago-Frangos, Dominick Faith, C. Martin Lawrence, Gabriel C. Lander, Blake Wiedenheft

**Author notes:** Current address: Department of Biochemistry and Cell Biology, Stony Brook University, Stony Brook, NY 11794. These authors contributed equally to this work (MaryClare F. Rollins, Saikat Chowdhury).

## Abstract

Bacteria and archaea have evolved sophisticated adaptive immune systems that rely on CRISPR RNA (crRNA)-guided detection and nuclease-mediated elimination of invading nucleic acids. Here we present the cryo-EM structure of the type I-F CRISPR RNA-guided surveillance complex (Csy complex) from *Pseudomonas aeruginosa* bound to a double-stranded DNA target. Comparison of this structure to previously determined structures of this complex reveals a *Ȉ*180-degree rotation of the C-terminal helical bundle on the “large” Cas8f subunit. We show that the dsDNA-induced conformational change in Cas8f exposes a Cas2/3 “nuclease recruitment helix” that is structurally homologous to a virally encoded anti-CRISPR protein (AcrIF3). Structural homology between Cas8f and AcrIF3 suggests that AcrIF3 is a mimic of the Cas8f “nuclease recruitment helix”, implying that cas genes may sometimes serve as genetic fodder for the evolution of anti-CRISPRs.

---

CRISPR (Clustered Regularly Interspaced Short Palindromic Repeats) and their associated genes (*cas*) are essential components of sophisticated adaptive immune systems that are widespread in bacteria and archaea, but are not found in eukaryotic genomes or in eukaryotic organelles that originated from bacteria (e.g., mitochondria and chloroplasts)^1-5^. Microbial CRISPR-Cas systems are divided into Class 1 systems, which rely on multi-subunit CRISPR RNA (crRNA)-guided surveillance complexes, and Class 2 systems, which rely on a single multi-domain protein that serves as a crRNA-guided effector nuclease^4,6^. The simple composition and programmable versatility of the Class 2 nucleases (i.e., Cas9, Cas12 and Cas13) has attracted considerable attention for diverse applications in genome engineering^7-9^. However, these systems are relatively rare in nature, occurring in fewer than 10% of sequenced bacterial and archaeal genomes, while the Class 1 systems represent the remaining 90% of adaptive immune systems observed in nature^6^.

Class 1 systems are divided into three different types (I, III, and IV) that are further divided into subtypes based on gene sequences and organization of the operon. The type I systems are the most abundant, widespread, and diverse of these systems, which include seven distinct subtypes (i.e., I-A through I-F; I-U)^4,6^. Despite this diversity, all type I systems rely on multi-subunit CRISPR RNA (crRNA)-guided surveillance systems to identify foreign DNA^10^, which is subsequently eliminated by the *trans*-acting nuclease-helicase, Cas3^11-18^. In most type I systems, Cas2 and Cas3 are separate proteins involved in adaptation (i.e., integration of foreign DNA into the CRISPR) and interference (i.e., crRNA-guided target degradation), respectively^6^. However, in I-F systems, these proteins are fused into a single polypeptide (i.e., Cas2/3) which forms a homodimer that assembles with four molecules of the Cas1 adaptation protein^19-21^. Within the Cas1-2/3 complex, the Cas1 proteins repress Cas2/3 endonuclease activity, which must be activated by the target bound type I-F surveillance complex (Csy complex)^20^. While previously determined structures of the Cas1-2/3 complex and the Csy surveillance complex provide mechanistic insights into their respective functions, the molecular mechanisms that govern Cas2/3 recruitment and nuclease activation remain obscure.

To understand the mechanism of target DNA recognition by the Csy surveillance complex, Guo *et al.* recently determined the structures of the Csy complex before DNA binding, and after binding to a partially duplexed DNA target^22^. These structures explain the mechanism of PAM recognition (Protospacer Adjacent Motif) and reveal an elongation of the complex that is driven by crRNA-guided hybridization to complementary DNA. However, the mechanism by which the nuclease is recruited to the target-bound complex was not elucidated. Here we use cryo-electron microscopy (cryo-EM) to determine the ∼3.2 Å-resolution structure of the Csy complex from *Pseudomonas aeruginosa* bound to an 80-basepair dsDNA target (Figure 1). The structure reveals dramatic conformational changes that are not observed in the previously determined structures. In combination with biochemical methods, we show that these structural differences have significant functional consequences. Specifically, this work explains how R-loop formation created by crRNA-guided strand invasion of a dsDNA target is necessary for driving a ∼180-degree rotation of the C-terminal helical bundle on the “large” Cas8f subunit. This conformational change presents a “nuclease recruitment helix” that is buried in the unbound structure. Additionally, we show that the helical bundle of Cas8f is structurally homologous to a virally-encoded anti-CRISPR protein (AcrIF3) that suppresses immune function by mimicking the nuclease recruitment helix on Cas8f, raising the possibility that *cas* genes may sometimes serve as genetic fodder for the evolution of anti-CRISPRs.

**Figure 1.**
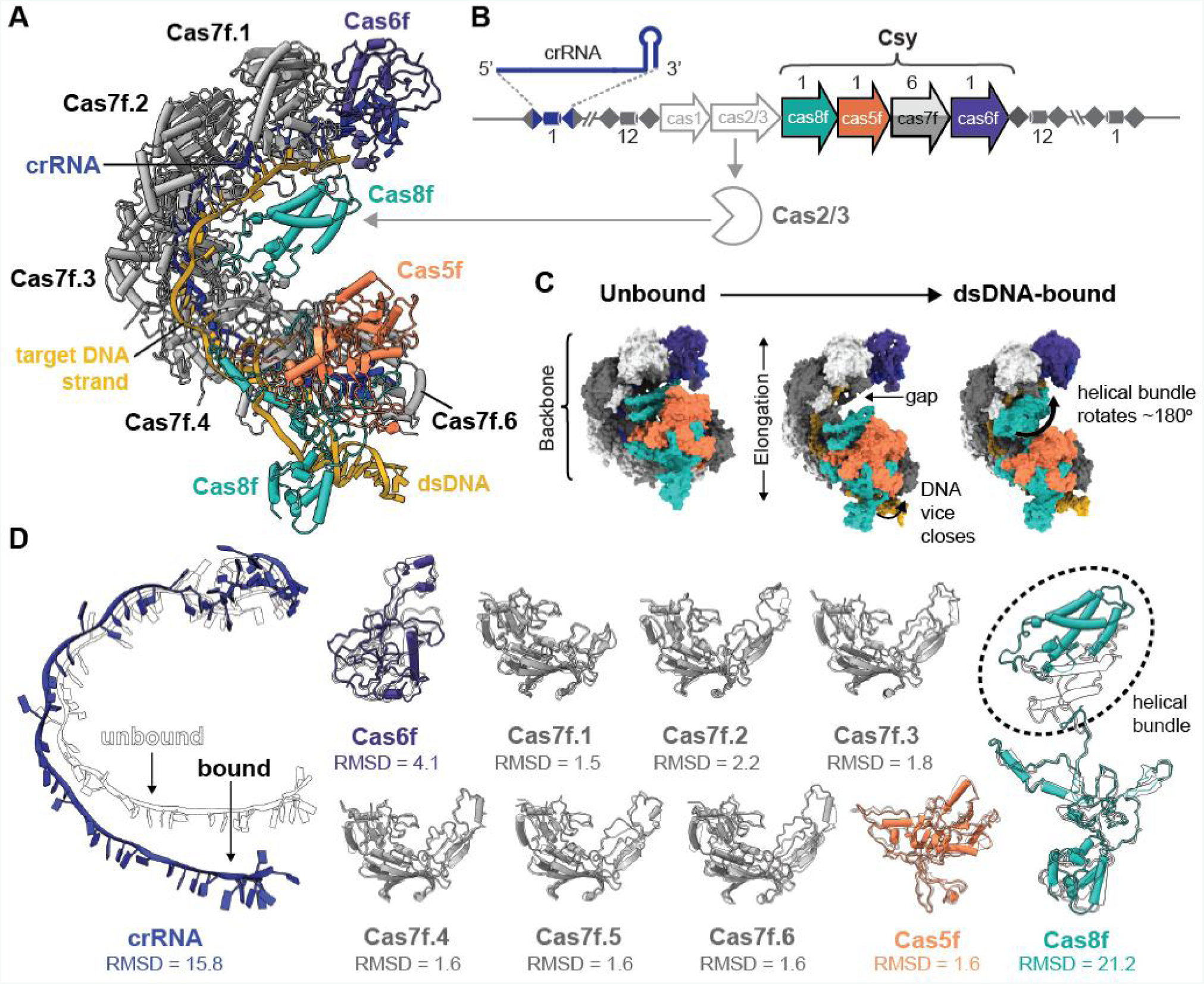
DNA binding induces conformational changes in the Csy complex. (A) Atomic model of the type I-F crRNA-guided surveillance complex (Csy complex) from *Pseudomonas aeruginosa* (PA14) bound to a dsDNA target. (B) The type I-F CRISPR-Cas immune system in *P. aeruginosa* (PA14) consists of six *cas* genes flanked by two CRISPR loci. Colored arrows indicate subunits within the Csy complex. The binding site for Cas2/3 (pac-man) is indicated with a gray arrow. (C) Schematic representation of the conformational change in the Csy complex, from unbound to dsDNA-bound. From L to R: the unbound complex (PDB ID: 6B45), Csy bound to a partially-duplexed dsDNA (PDB ID: 6B44), and the dsDNA-bound complex (PDB ID: 6MPU).

## DNA binding induces conformational changes in the Csy complex

To determine the mechanism of foreign DNA recognition and Cas2/3 recruitment by the Csy complex, we determined the ∼3.2 Å cryo-EM structure of the Csy complex from *Pseudomonas aeruginosa* (strain PA14) bound to an 80-basepair dsDNA target containing a protospacer and a PAM (Figure 1A, S1-S3, Table S1-S2). The cryo-EM reconstruction was of sufficient quality for atomic modeling (see methods). The 3’ prime end of the R-loop and a fifteen-residue linker within the Cas8f subunit could not be modeled due to lack of density in the reconstructed map, which may be due to intrinsic flexibility.

The Csy complex is a multi-subunit crRNA-guided surveillance complex composed of an unequal stoichiometry of four different CRISPR-associated (Cas) proteins, and a single 60-nt crRNA (Cas8f_1_:Cas5f_1_:Cas7f_6_:Cas6f_1_:crRNA_1_)^22-25^ (Figure 1B). The complex assembles into an asymmetric spiral that is capped at one end by Cas6f (i.e., the “head”) and on the other by a heterodimer of Cas5f and Cas8f, which form the “tail”. Cas6f (formerly Csy4) is a CRISPR RNA processing enzyme that binds to and cleaves CRISPR RNA stem-loop structures consisting of palindromic repeat sequences^26-28^. After cleavage, Cas6f remains stably associated with the 3’ end of the crRNA, and six Cas7f subunits oligomerize along the crRNA, forming the “backbone” of the complex^22,24,25^ (Figure 1A). In the tail, the 5’ end of the crRNA is anchored by a network of interactions within the stable heterodimer formed by Cas5f and the N-terminal domain of Cas8f.

The dsDNA target-bound structure undergoes significant conformational rearrangements relative to both the unbound complex and the complex bound to a partial duplex^22^ (Figure 1C, Movie S1), while retaining the same overall morphology (head, backbone, and tail). The transition to the dsDNA-bound conformation can be broadly described in three coordinated movements. First, a positively-charged “DNA vise” formed by the N-terminal segment of Cas8f and the opposing face of Cas7f.6 closes around the dsDNA. In this position, two loops of Cas8f insert into the DNA minor groove, where specific residues interact with the Protospacer Adjacent Motif (PAM). Cas8f and Cas5f form a stable heterodimer^23,24^ and movement of the N-terminus of Cas8f coincides with a ∼25 Å rigid-body translation of Cas5f away from the head of the complex. This action, combined with hybridization between the target DNA and the complementary crRNA spacer, results in an elongation of the Cas7f backbone. Compared to the unbound conformation, the length of the backbone as measured from Cas7f.1 to Cas7f.6 is extended ∼18 Å in the target-bound structure, which opens the tight helical spiral, exposing an average of ∼145 Å^2^ of formerly buried surface area between adjacent Cas7 subunits. The elongated conformation also creates a gap between the head and the tail of the complex that is necessary for a ∼180-degree rotation of the helical bundle of Cas8f (Figure 1C).

Transition to the dsDNA-bound conformation is primarily accomplished by rigid-body rearrangements of the Cas subunits; structures of individual subunits reveal few changes relative to their unbound state (Figure 1D). Notably, the first two conformational changes (i.e., closing of the DNA vice, and elongated Cas7f backbone) are evident in a recent cryo-EM structure of the Csy complex bound to a partial dsDNA target^22^. However, rotation of the Cas8f helical bundle is absent in this structure, suggesting that this rearrangement is dependent on R-loop formation. The dsDNA-bound structure presented here also reveals a “locked” conformation not observed in previous models, where regions of Cas7f.2 and Cas7f.3 fold over the DNA target strand and contact the helical bundle of Cas8f, completely encasing the complementary DNA. Thus, target binding triggers dramatic conformational changes in the Csy complex, and some of these rearrangements are only observed when Csy binds a fully duplexed DNA target.

## Cas8f mediates dsDNA binding and PAM recognition

In type I systems, crRNA-guided surveillance complex initially engages DNA through non-sequence-specific electrostatic interactions with dsDNA, followed by sequence-specific interactions with the protospacer adjacent motif (PAM)^29-32^. PAMs (protospacer adjacent motifs) are short sequence motifs that flank the protospacer in foreign targets only, thereby distinguishing self-DNA from non-self-DNA^33,34^ (Figure 2A). PAM recognition by the surveillance complex destabilizes the DNA duplex and facilitates crRNA-guided strand invasion^22,35,36^. Hybridization of the crRNA-guide to the complementary DNA displaces the non-complementary strand, resulting in an R-loop structure^35-42^. The N-terminal domain of Cas8f and the opposing face of the terminal Cas7f subunit (Cas7f.6) form a positively charged “vise” that closes around dsDNA (Figure 2B-C)^22,24^. DNA binding results in a conformational change that moves the positively charged arm of Cas8f (R24-R58) ∼15Å into the closed position, clamping the complex onto the dsDNA (Figure 2C). In addition, closing of the DNA vise positions two loops of Cas8f in the DNA minor groove, which coincides with local distortion of the helix and separation of the DNA strands (Figure 2D-F, Movie2 S1,S2). Asparagine 111 (N111) and asparagine 250 (N250) of Cas8f are positioned within hydrogen bonding distance of the −2 and −1 positions of the PAM, respectively (Figure 1D). To verify the role of these residues in PAM recognition, we introduced alanine mutations at N111 and N250. While we were unable to purify the Csy complex containing the N111A mutation in Cas8f, the N250A mutant expressed and purified like wild-type (WT) complex (Figure S4). We performed electrophoretic mobility shift assays (EMSAs) with both WT and Cas8f N250A Csy complex (Figure 2G, Figure S4). Compared to WT, the Cas8f N250A mutation decreased DNA binding affinities by ¿3 orders of magnitude.

**Figure 2:**
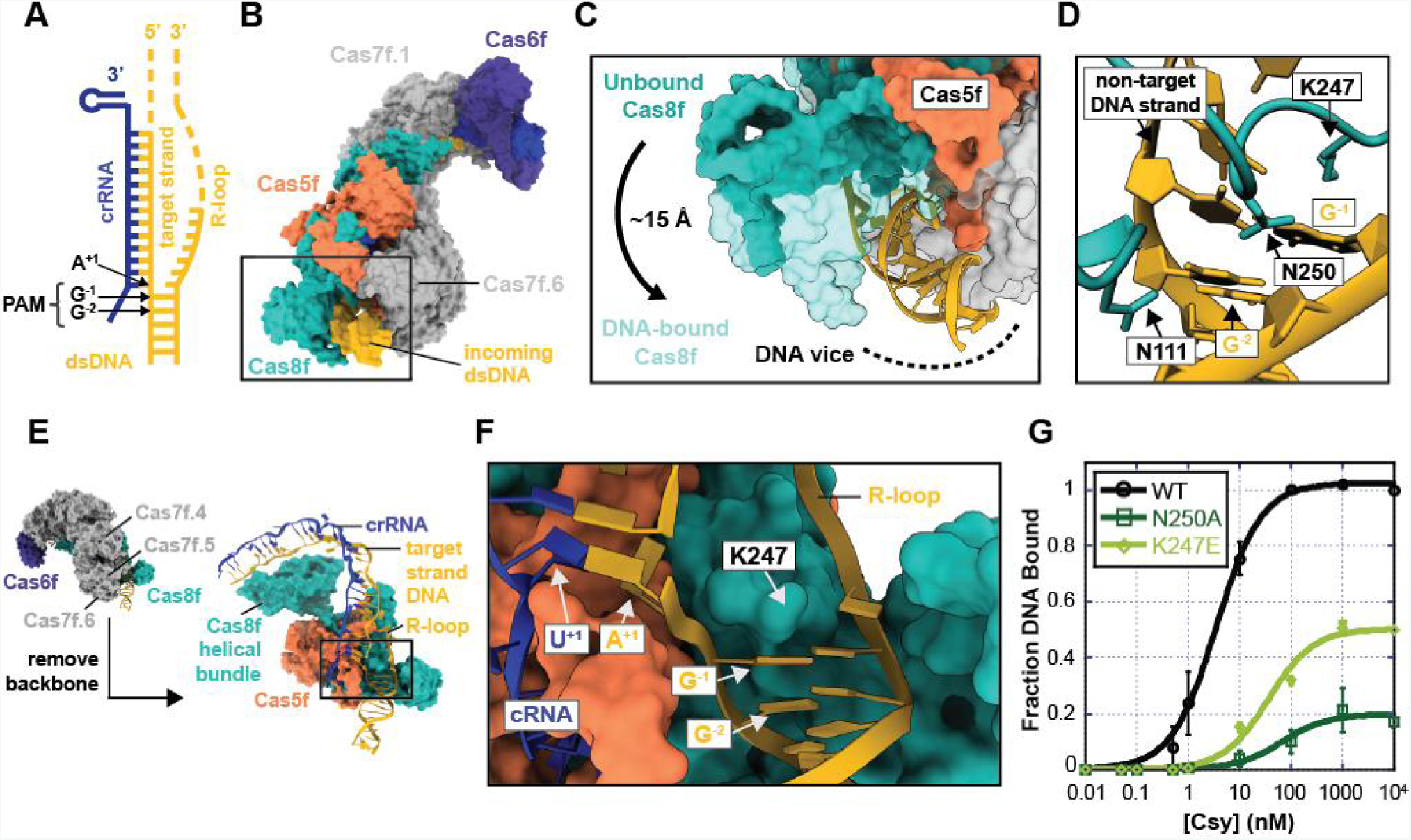
Cas8f and Cas7.6 form a vise that closes on dsDNA and recognizes the PAM. (A) Schematic of 80-nucleotide dsDNA target bound by the Csy complex. Dashed segments of the DNA (yellow), represent regions of the target that were not sufficiently ordered and could not be reliably modeled. (B) Surface representation of the dsDNA-bound Csy complex. The “DNA vise” (black box) is formed by the N-terminal domain of Cas8f and the opposing face of Cas7.6f. (C) Conformational change of the vise upon dsDNA binding. The positively charged arm of Cas8f (R24-R58) moves ∼15 Å into the closed position. (D) Two loops of Cas8f are inserted into the minor groove. Asparagine 111 (N111) is positioned within hydrogen bonding distance of the second base-pair of the PAM (i.e., G-C^-2^), and asparagine 250 (N250) is oriented toward the −1 G of the PAM (G^-1^). (E) Sidelong view of the dsDNA-bound Csy complex. The box highlights Cas8f-mediated DNA strand splitting. (F) Lysine 247 (K247) acts as a wedge, separating the strands and positioning the first nucleotide of the target sequence for base-pairing with the first nucleotide of the crRNA guide. (G) Electrophoretic mobility shift assays performed with radiolabeled dsDNA substrates show that Cas8f mutations N250A or K247A result in reduced crRNA-guided DNA binding. Error bars, SD; n = 3.

The DNA strands separate at the first base-pair of the protospacer (i.e. position +1). Strand-splitting is facilitated by lysine 247 (K247), which forms a wedge that inserts between the strands above the PAM (Figure 2D-F). To test the requirement of this wedge for target binding, we introduced a charge-swap mutation (K247E) in Cas8f and measured its impact using EMSAs. The K247E mutation results in a binding defect and corroborates previous structural observations of the Csy complex bound to a partially-duplexed DNA target (Figure 2G)^22^. In fact, comparison of the two structures suggests the mechanism of PAM recognition is unchanged by the presence or absence of an R-loop. The root-mean-square deviation (RMSD) for equivalently positioned C-alpha atoms in the Cas8f NTDs is 1.69 Å. This is consistent with an early role for PAM recognition in target binding, prior to formation and coordination of the R-loop.

## The interface between Cas8f and Cas5f forms an R-loop binding channel

PAM recognition induces local distortion of the DNA duplex that facilitates crRNA-guided hybridization to the complementary DNA target, which displaces the non-complementary DNA strand (R-loop). The first nine-nucleosides of the R-loop are positioned along a positively charged channel formed by residues in Cas8f (K28, K31, K119, R207, R219, R258, R259) and Cas5f (K76, R77) (Figure 3A, Movie S2). While the density for the remaining nucleosides of the flexible R-loop are insufficiently resolved for atomic modeling, the positively charged channel continues along Cas5f and the helical bundle of Cas8f, terminating between arginine-rich helixes on Cas5f (K171, R175, R178, R179) and Cas8f (R293, R299, R302, R306) (Figure 3B). We hypothesized that this positively charged channel stabilizes the DNA-bound conformation by binding the R-loop and limiting reannealing of the DNA duplex. To test this hypothesis, we introduced positive-to-negative charge-swap mutations along the length of the channel. A quadruple mutant (R282E/R293D/R299E/R302E) of residues in the helical bundle of Cas8f expressed and purified similar to WT Csy complex (Figure S4). Mutations to the “R-loop binding channel” (RBC) result in a substantial dsDNA binding defect (Figure 3C and S4). To confirm that this binding defect is a function of decreased R-loop stability, we repeated the experiment with a dsDNA substrate containing a non-complementary “bubble”, which would form an R-loop incapable of reannealing. Consistent with our hypothesis, the RBC mutant bound the DNA bubble with WT binding affinity, demonstrating that the positive charge in this channel plays an important role in R-loop stabilization. Together, our structural and biochemical data suggest the RBC makes sequence-independent interactions with the R-loop that inhibit reannealing of the DNA duplex.

**Figure 3:**
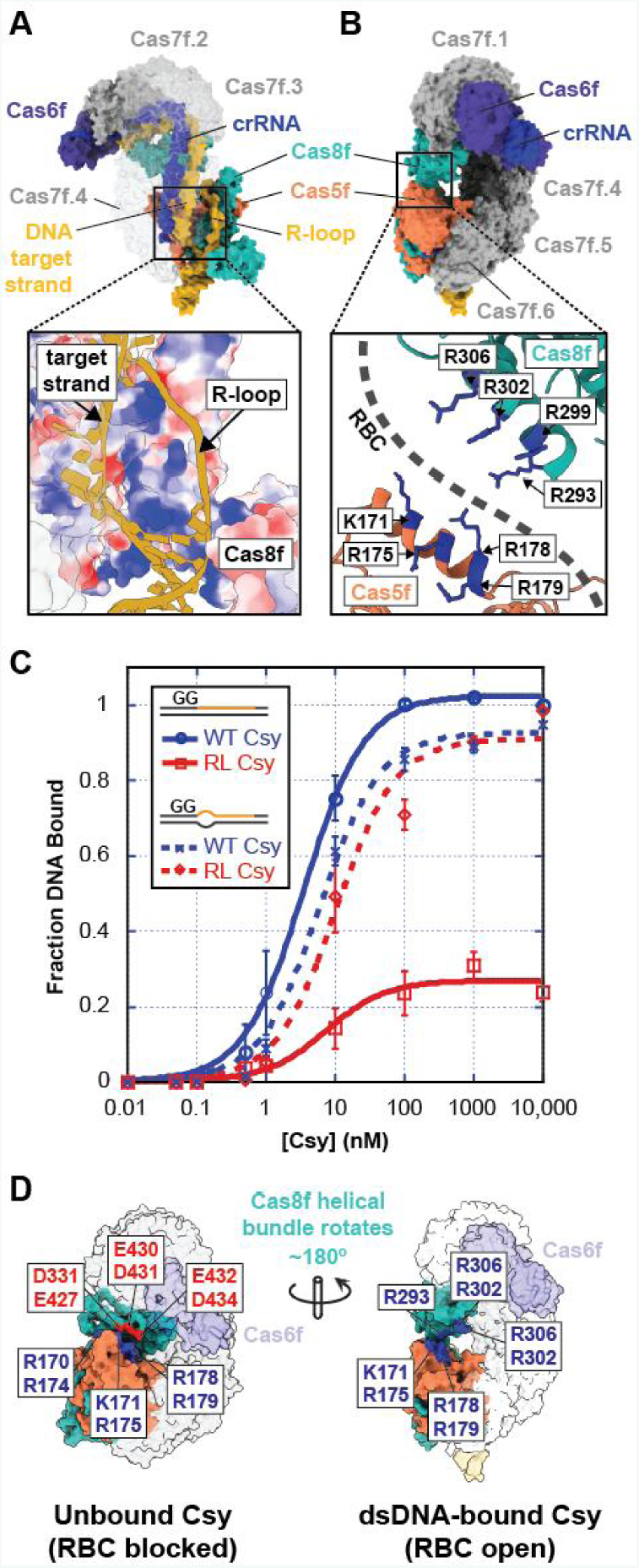
The non-complementary strand is positioned in a positively charged channel. (A) Surface representation of the dsDNA-bound Csy complex, with inset showing the non-complementary strand (R-loop) positioned in a positively-charged (blue) channel formed by residues in Cas8f and Cas5f. (B) Ninety-degree rotation of the DNA-bound Csy complex. Inset shows the PAM-distal end of the R-loop binding channel (RBC), formed by arginine-rich helices on Cas5f and Cas8f. (C) Electrophoretic mobility shift assays performed with ^32^ P-labeled dsDNA substrates show that charge-swap mutations in Cas8f residues R282/R293/R299/R302 result in reduced dsDNA binding. However, high-affinity binding is rescued by DNA targets with 10-nucleotide protospacer “bubbles”. Error bars, SD; n = 3 (D) Positions of the Cas8f and Cas5f RBC helices in unbound and target-bound Csy. In unbound Csy, the Cas8f RBC helix is positioned on the interior of the complex and the Cas5f RBC helix is occupied by shape and charge complementation with acidic residues on Cas8f (D331, E427, E430, D431, E432, D434). In target-bound Csy, the Cas8f helical bundle is rotated, completing the RBC.

The PAM-distal end of the RBC is composed of arginine-rich helices on Cas5f and the helical bundle of Cas8f (Figure 3B). Notably, formation of this section of the RBC requires rotation of the Cas8f helical bundle, and rotation of the helical bundle requires dsDNA binding. When the Csy complex is unbound, the helical bundle of Cas8f is not rotated, and the Cas5f RBC helix (D166-R179) is juxtaposed with acidic residues on Cas8f (D331, E427, E430, D431, E432, D434) (Figure 3D). In fact, this interaction between Cas5f and the unrotated Cas8f helical bundle is preserved in a structure of the Csy complex bound to dsDNA with an incomplete R-loop^22^. This observation suggests R-loop binding along the length of the RBC may disrupt the charge-complementation between Cas8f and Cas5f, allowing for rotation of the helical bundle.

## The R-loop is a regulator of Cas2/3 recruitment

Type I-F CRISPR defense is initiated when the Csy complex binds a dsDNA target, which leads to recruitment of the *trans*acting nuclease/helicase Cas*2*/3 for DNA degradation^20,29,43-46^. However, Cas2/3 forms a complex with the adaptation protein Cas1, and Cas1 inhibits Cas2/3 nuclease activity^20,21,44^. Because the Cas2/3 nuclease is activated by the DNA-bound Csy complex, we reasoned that the recruitment signal must be coincident with the conformational change that occurs during dsDNA binding. To test this hypothesis, we performed ESMAs with purified Csy complex, purified Cas1-2/3 complex, and ^[32]^ P-labeled dsDNAs designed to determine how specific features of the DNA ligand impact Cas2/3 recruitment. First, we measured Cas1-2/3 recruitment to Csy complex bound to a dsDNA target with a full protospacer and a GC-GC PAM, using electrophoretic mobility shift assays (EMSAs) (Figure 4A). As previously reported, Cas1-2/3 recruitment results in two supercomplexes. The lower molecular weight complex contains dsDNA, Csy and Cas2/3, while the larger, more transient complex that may include Cas1 (i.e., dsDNA, Csy and Cas1-2/3)^20^. As expected, increasing concentrations of Cas1-2/3 complex resulted in loss of the band corresponding to the dsDNA-bound Csy complex and a corresponding increase in the intensity of the bands representing dsDNA-Csy-Cas2/3 supercomplexes (Figure 4C and S5A). Next, we tested Cas2/3 recruitment to Csy complex bound to a partially-duplexed target like the one used by Guo *et al.*^22^, whose structure contained a closed DNA vise and an elongated Cas7f backbone, but did not show a rotation of the helical bundle (Figure 4B). We hypothesized that Csy bound to this partially duplexed substrate would be unable to recruit Cas2/3, as the orientation of the Cas8f helical bundle would prevent access to the necessary docking site. Indeed, results of the EMSA experiments indicate that the partial DNA duplex does not support recruitment of the nuclease (Figure 4C). These results suggest that the R-loop is necessary for stable rotation of the helical bundle, and that rotation of the helical bundle is critical for stable association with Cas2/3.

**Figure 4:**
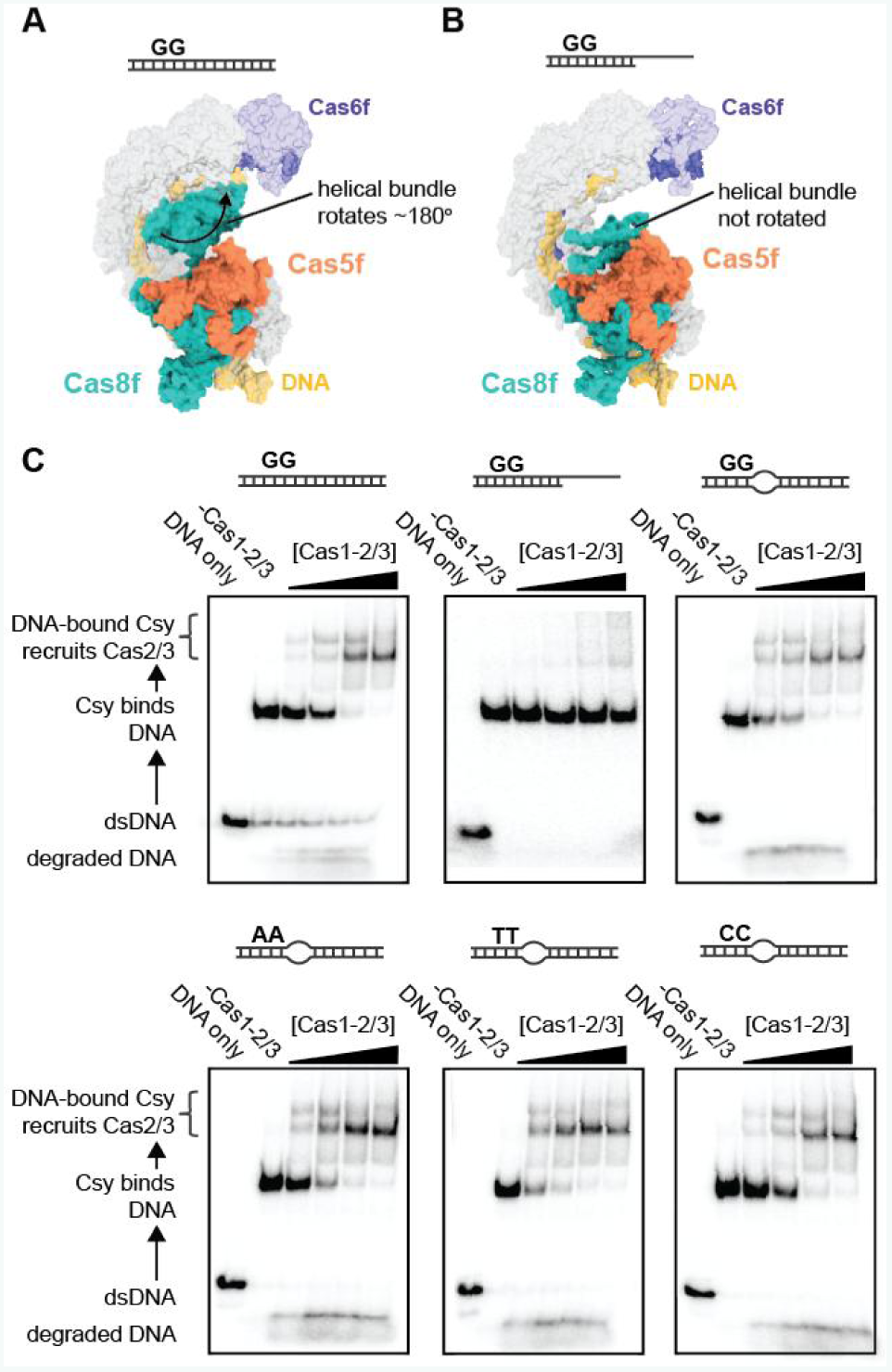
The R-loop is a regulator of Cas2/3 recruitment. (A) Model of the Csy complex bound to a complete dsDNA target (schematic included above). The Cas8f helical bundle is rotated ∼180° relative to the unbound conformation. (B) Model of the Csy complex bound to a partial dsDNA target (schematic included above) (PDB: 6B44). The Cas8f helical bundle is not rotated relative to the unbound conformation. (C) Electrophoretic mobility shift assays (EMSAs) were performed with radiolabeled dsDNA substrates (illustrated schematically above each gel), purified Csy complex and increasing concentrations (1.85 nM, 5.5 nM, 16.6 nM or 50 nM) of the Cas1-2/3 complex. Quantification of EMSAs (Figure S5A) show a Cas1-2/3-dependent decrease in dsDNA-bound Csy complex, and corresponding increase in dsDNA-Csy-Cas2/3 supercomplex. This was seen for all DNA substrates tested except the partial duplex.

There is evidence that the PAM serves as an allosteric regulator of Cas3 recruitment in type I-E systems^31,47,48^. To test whether the PAM regulates Cas2/3 recruitment to the Csy complex, we performed EMSAs with targets containing a canonical double-stranded G-C/G-C PAM or a T-A/T-A, AT/A-T, or C-G/C-G PAM (Figure 4C). The Csy complex has a stringent requirement for a PAM composed of two consecutive G-C base pairs, and PAM mutations result in severe DNA binding defects^29^. To facilitate binding to DNA targets with mutant PAMs, we used dsDNA targets with a 10-nt bubble in the protospacer (positions 1 - 10) (Table S1). PAM mutants that contain the 10-nt bubble are bound with near-WT affinities but unlike what has been reported in type I-E systems, the mutant PAMs had no effect on Cas2/3 recruitment or DNA degradation, suggesting that the PAM is necessary for crRNA-guided strand invasion of the DNA duplex, but the PAM does not regulate Cas2/3 activity (Figure 4C).

## Target-bound Csy complex assumes a “locked” conformation

In addition to its role in Cas2/3 recruitment, rotation of the Cas8f helical bundle may contribute to the stable “locked” conformation of the dsDNA-bound Csy complex. The Csy complex stably associates with dsDNA targets that include a PAM and a complementary protospacer (*K*_D_∼1 nM)^24,29^. This binding behavior is similar to what has been reported for DNA binding by the type I-E surveillance complex (i.e., Cascade). In I-E systems, target-bound Cascade assumes a locked conformation, resulting in an extended half-life on DNA targets. Locking involves the translocation of two subunits (Cse2 proteins) that pinch the DNA target during binding^35-40,48^. While the type I-E and I-F surveillance complexes share morphological similarities, the I-F complex does not contain Cse2 homologs. Instead, the dsDNA-bound structure of Csy reveals an alternative locking mechanism that involves two of the six Cas7f backbone subunits (Figure 5A).

**Figure 5:**
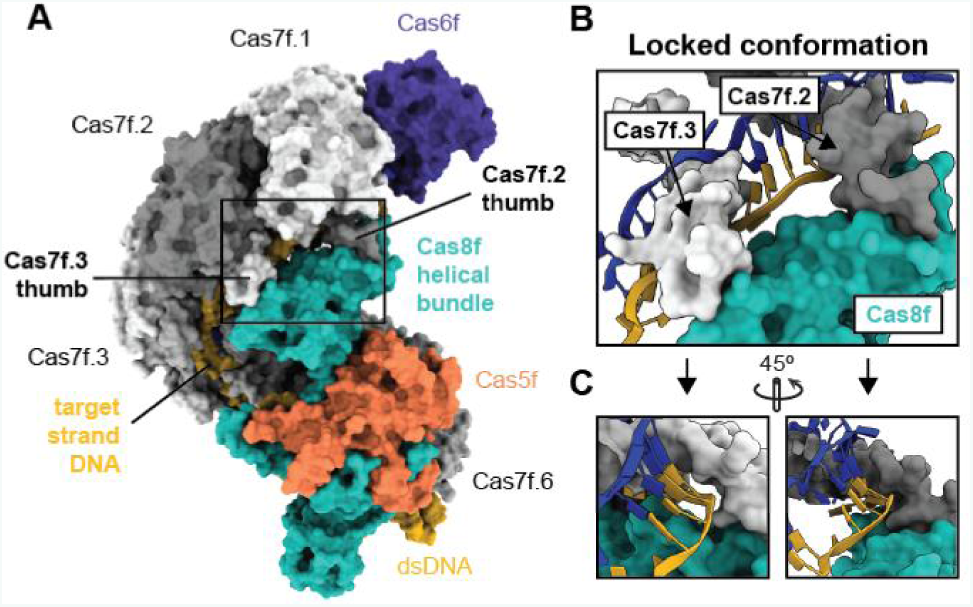
Target-bound Csy complex adopts a locked conformation. (A) Surface model representation of dsDNA-bound Csy complex. The target DNA strand is encapsulated by contacts between the helical bundle of Cas8f and the thumbs of Cas7f.2 and Cas7f.3. (B) Detail of the locked conformation showing the thumbs of Cas7f.2 and Cas7f.3 (T71-N94) piercing the crRNA-DNA duplex, then folding over the top of the complementary strand and interacting with Cas8f. (C) The complementary DNA strand is completely encased by the Cas7f thumbs and the helical bundle of Cas8f.

Like all other Cas7 family proteins, Cas7f proteins have a characteristic “right-hand” morphology composed of fingers-, palm-, web-, and thumb-shaped domains^24^. Each of these proteins “grip” the crRNA though non-sequence specific interactions with the phosphate backbone via residues distributed across each of the Cas7f domains. The thumb folds over the crRNA at regular six-nucleotide intervals in a way that precludes base-pairing at each of these positions. Thus, hybridization between the crRNA and the complementary DNA results in five-base pair segments of duplex that are interrupted at every sixth position by a thumb. The importance of the thumb in partitioning the crRNA into discrete segments has been well-established, but the structure of the dsDNA-bound complex reveals that, after piercing the crRNA-DNA duplex, the thumbs of Cas7f.2 and Cas7f.3 (T71-N94) curl over the top of the complementary strand and interact with the helical-bundle on Cas8f, creating a tunnel that fully encircles the complementary strand of DNA (Figure 5B). This structural conformation appears to lock the complex in a DNA-bound state and may explain the extended half-life of the target-bound Csy complex.

## Anti-CRISPR AcrIF3 is a molecular mimic

Bacteriophages (phages) have evolved numerous mechanisms to subvert CRISPR defense^49-51^. Several temperate phages of *P. aeruginosa* encode small proteins that bind and neutralize type I-F Cas proteins^22,24,51-58^. One of these anti-CRISPR proteins (AcrIF3) binds Cas2/3 and prevents its recruitment to the Csy complex^20,52,54,55^. The structure of AcrIF3 is similar to the helical bundle of Cas8f, suggesting that this anti-CRISPR may function as a molecular mimic^24^ (Figure 6A). When we compared structures of the two proteins, we identified one helix with conserved amino acids (Figure 6B-C). Crystal structures of Cas2/3 bound by AcrIF3 indicate that conserved residues on AcrIF3 form a hydrogen-bonding network with the C-terminal domain (CTD) of Cas2/3, and mutations in these residues abolish AcrIF3 binding^54,55^. We wondered whether the corresponding residues on the Cas8f helical bundle were binding Cas2/3 in a similar way.

**Figure 6:**
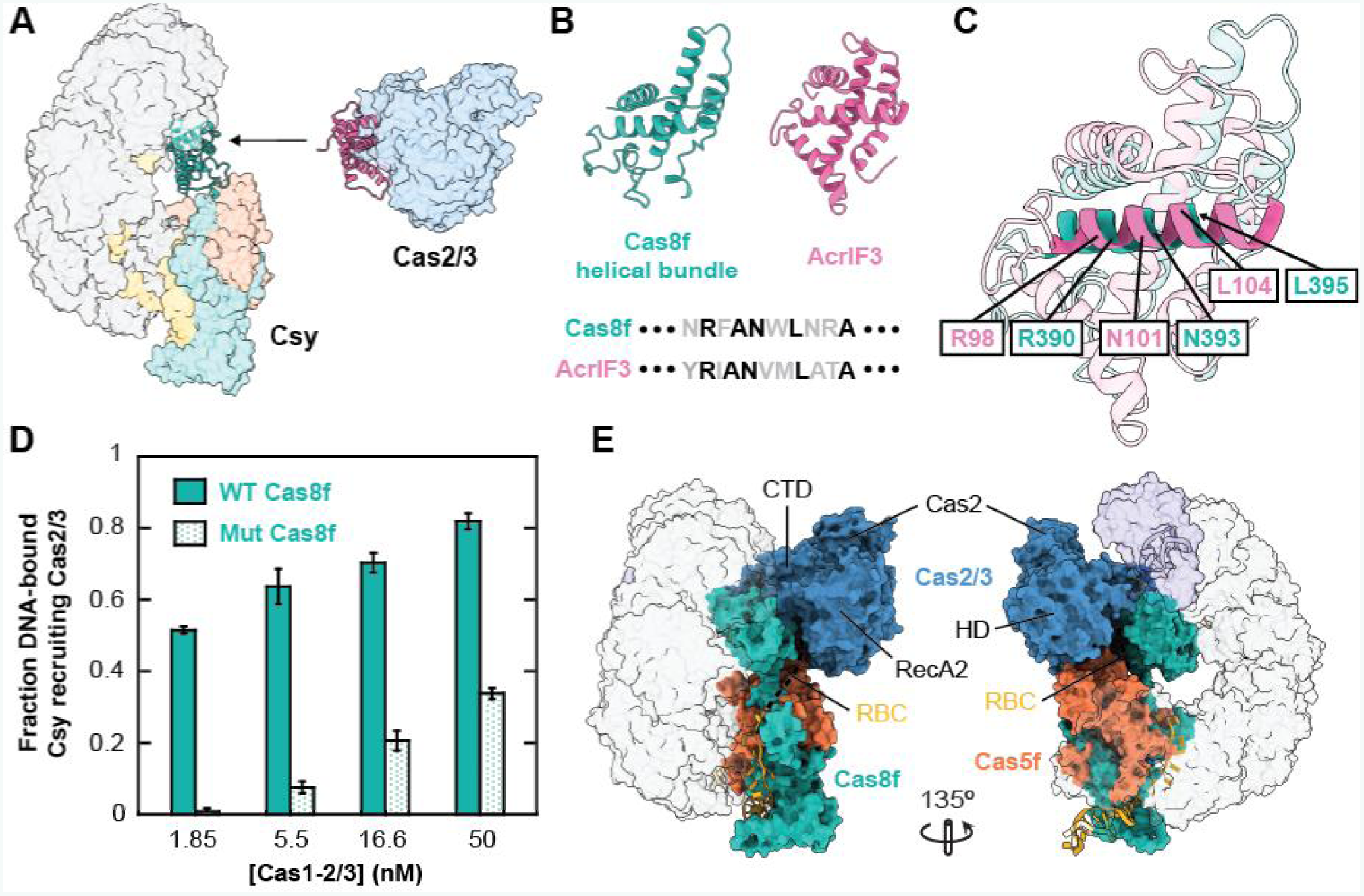
Anti-CRISPR mimicry reveals Cas2/3 docking site on Csy. (A) Models of target-bound Csy complex (left) and Cas2/3 bound by the anti-CRISPR AcrIF3 (right). AcrIF3 (pink) and the helical bundle of Cas8f (green) are shown as ribbons. (B) Structures of the Cas8f helical bundle and phage-encoded anti-CRISPR protein AcrIF3 and amino acid sequence conservation between the two proteins. (C) Alignment of Cas8f helical bundle (green) and AcrIF3 (pink) with conserved helix in foreground. Positions of mutated residues are indicated. (D) Electrophoretic mobility shift assays (EMSAs) were performed with radiolabeled dsDNA, purified Csy complex and 1.85 nM, 5.5 nM, 16.6 nM or 50 nM Cas1-2/3 complex. Quantification of Cas2/3 recruitment shows a defect in the Cas8f R390A and N393A mutants relative to WT. Error bars, SD; n = 3. (E) Model of dsDNA-Csy-Cas2/3 supercomplex. Cas2/3 was docked on to dsDNA-bound Csy by aligning AcrIF3 with the Cas8f helical bundle.

To test this hypothesis, we made alanine point mutations in conserved residues R390, N393, and L395 (Figure 6B-C). The mutations result in a Cas2/3 recruitment defect (Figure 6D and S5B). This result also supports our hypothesis that AcrIF3 blocks CRISPR defense by mimicking the helical bundle of Cas8f and occupying its binding site on Cas2/3. We took advantage of this mimicry to generate a model of the DNA-Csy-Cas2/3 supercomplex. We aligned AcrIF3 with the rotated helical bundle of Cas8f to dock Cas2/3 onto the target-bound Csy complex (Figure 6E). In the resulting model, Cas2/3 contacts the Cas8f helical bundle and parts of the N-terminal region of Cas5f. In this position, the R-loop binding channel (RBC) directs the displaced DNA strand into the RecA helicase domains of Cas2/3. The location of the Cas2/3 HD nuclease domain near the end of the R-loop is also consistent with previous data indicating Cas2/3 initially nicks the R-loop in a PAM-distal position^20^. We expect this model will help direct further investigation of Cas2/3 recruitment and supercomplex formation in the type I-F system.

## Discussion

Here we describe the mechanism by which a type I-F crRNA-guided surveillance complex recognizes dsDNA and signals recruitment of the Cas2/3 nuclease-helicase to degrade a *bona fide* target. We determined the cryo-EM structure of the type I-F crRNA-guided surveillance complex from *P. aeruginosa* bound to a dsDNA target (Figure 1), and compared it to a recently published structure of the complex bound to a partially duplexed DNA^22^. Surprisingly, we observe a major conformational difference that requires binding dsDNA, which is the biologically relevant target. We show that both the complementary and non-complementary strands of the target duplex have distinct but coordinated roles in transitioning the complex into a nuclease-ready conformation. Hybridization between the crRNA-guide and the complementary DNA is necessary for elongation of the Cas7f backbone, while displacement of the non-complementary strand (i.e. R-loop formation) is directly involved in rotating the C-terminal helical bundle of Cas8f. These two rearrangements (i.e., elongation and rotation) are coordinated by directional unwinding of the duplex. Rotation of the Cas8f helical bundle creates a positively charged groove between Cas8f and Cas5f that stabilizes the R-loop and inhibits reannealing of the DNA duplex (Figure 3). The rotated conformation of Cas8f is stabilized by the “thumbs” of Cas7f.2 and Cas7f.3, which fold over the complementary DNA target and contact the helical bundle of Cas8f, completely encasing the complementary DNA target (Figure 5). This conformation provides a structural explanation for the extended half-life of the Csy complex on a DNA target, and also indicates that this stabilized, or “locked” configuration can only occur after base pairing extends to the 3’-end of the crRNA guide. This locking process is conceptually similar to locking mechanisms that have been described for the type I-E systems and conformational control mechanisms that have been reported for Cas9^37,59-61^.

While the coordinated movements of the Csy surveillance complex serve as a dynamic example of conformational versatility (Movie S1), the biological function of the observed conformational rearrangements were not immediately evident. In particular, it was unclear if the ∼180-degree rotation of the Cas8f helical bundle had functional importance beyond the locking process describe above. We previously identified structural homology between the anti-CRISPR protein AcrIF3 and this helical bundle^24^, and given that AcrIF3 binds Cas2/3^52,54,55^, we hypothesized that the helical bundle may similarly interact with Cas2/3. To test this hypothesis, we initially superimposed structures of AcrIF3 bound to Cas2/3 onto the helical bundle of Cas8f. Performing this superposition using structures of the Csy complex before DNA binding or after binding to a partially duplex DNA resulted in substantial steric clashes between Cas2/3 and the Cas7f backbone (Figure 7, Movie S3). However, the structure presented here shows that dsDNA binding re-orients the helical bundle into a position that can accommodate Cas2/3 binding, aligning structural features of Cas2/3 with complementary features on Cas8f and Cas5f. Moreover, the position of the Cas2/3 nuclease domain is consistent with previous biochemical data suggesting that cleavage of the R-loop occurs at the PAM-distal end of the protospacer^20^.

**Figure 7:**
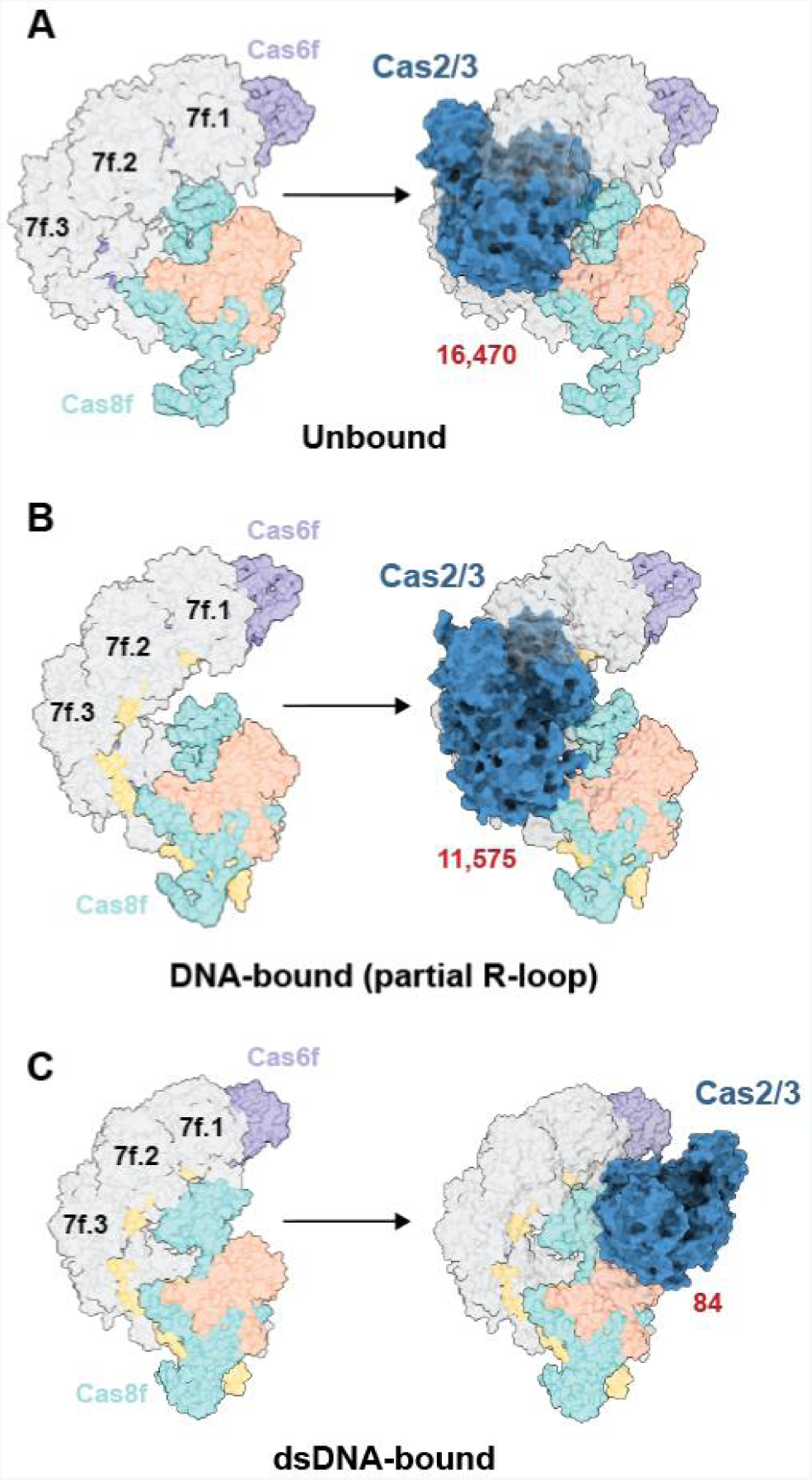
Double-stranded DNA-induced conformational change in Cas8f exposes a Cas2/3 “nuclease recruitment helix”. (A-C) Surface models of the Csy complex (unbound Csy PDB ID: 6B45; Csy bound to partially-duplexed DNA PDB ID: 6B44; dsDNA-bound Csy PDB ID: 6MPU). Cas2/3 (blue) was docked onto each model by aligning AcrIF3 with the Cas8f helical bundle. Red numbers indicate the number of clashing atoms between Cas2/3 and Csy.

Collectively, our structural and biochemical analyses not only revealed a mechanistic model for nuclease recruitment to a CRISPR RNA-guided surveillance complex, but also demonstrates how the anti-CRISPR protein AcrIF3 subverts type I-F CRISPR defense through molecular mimicry. While numerous anti-CRISPRs have now been shown to function as mimics of DNA^22,24,62-64^, this is the first example of an anti-CRISPR that mimics a Cas protein, and suggests that *cas* genes themselves may serve as genetic fodder for the evolution of anti-CRISPR proteins. This study emphasizes the importance of anti-CRISPRs as tools to understand the functions of CRISPR-Cas systems they target.

## Materials and Methods

### Protein expression and purification

#### P. aeruginosa Csy complex

Csy genes and a synthetic CRISPR were co-expressed on separate vectors in *E. coli* BL21 (DE3) cells as previously described^20^. Expression was induced with 0.5 mM isopropyl-D-1-thiogalactopyranoside (IPTG) at an optical density (OD 600 nm) ∼0.5. Cells were incubated overnight at 16°C, then pelleted by centrifugation (5000 × g for 15 min at 4°C) and re-suspended in lysis buffer (50 mM 4-(2-hydroxyethyl)-1-piperazineethanesulfonic acid (HEPES) pH 7.5, 300 mM potassium chloride, 5% glycerol, 1 mM Tris(2-carboxyethyl) phosphine hydrochloride (TCEP), 1x protease inhibitor cocktail (Thermo Scientific)). Pellets were sonicated on ice for 3 × 2.5 min (1 sec on, 3 sec off), then lysate was clarified by centrifugation at 22,000 × g for 30 min at 4°C. The Csy complex self-assembles *in vivo* and the intact complex (with N-terminal 6-histidine affinity tags on Cas7f) was affinity purified over NiNTA resin (Qiagen) which was washed once with lysis buffer supplemented with 20 mM imidazole before elution with lysis buffer supplemented with 300 mM imidazole. Protein was then concentrated (Corning Spin-X concentrators) at 4°C before further purification over a Superdex 200 size-exclusion column (GE Healthcare) in 20 mM HEPES pH 7.5, 100 mM KCl, 5% glycerol, 1 mM TCEP.

#### P. aeruginosa Cas1-2/3 complex

The Cas1-2/3 complex was expressed and purified using previously described methods and the plasmids are available on Addgene (#89240)^20^. Briefly, the expression vector was transformed into *E. coli* BL21 (DE3) cells, and the cells were induced with IPTG at an OD_600_ of 0.5. Expression was induced with 0.5 mM IPTG at OD_600_ = 0.5 nm. Cells were pelleted and lysed as described above. Co-expressed Cas1 (with N-terminal 6-histidine affinity tag) and Cas2/3 (untagged) were affinity purified using NiNTA resin (Qiagen), which was washed once with lysis buffer supplemented with 20 mM imidazole before elution with lysis buffer supplemented with 300 mM imidazole. Protein was concentrated (Corning Spin-X concentrators) at 4°C before further purification over a Superdex 200 size-exclusion column (GE Healthcare) in 20 mM HEPES pH 7.5, 100 mM KCl, 5% glycerol.

### Grid preparation for cryo-electron microscopy

Prior cryo-EM studies with the Csy-Acr complex^24^ showed that Csy complexes adopt a preferred orientation in ice. Addition of 0.05% (v/v) Lauryl Maltose Neopentyl Glycol (LMNG, Anatrace) to the sample helped in overcoming this orientation bias problem. 4 *µ*L of 2 mg/mL purified Csy-DNA complex, mixed with 0.05% (v/v) LMNG was added onto freshly plasma cleaned (hydrogen, oxygen plasma) 300 mesh UltrA-uFoil R1.2/1.3 holey Gold grid (Quantifoil). After manually blotting off excess sample with a Whatman No.1 filter paper for 5-7 s, the sample was immediately vitrified by plunge freezing in liquid-ethane at −179°C. The entire cryo grid preparation process was carried out at 4°C and 98% relative humidity to minimize excessive evaporation of sample from grid surface.

### Cryo-electron microscopy data acquisition

Cryo grids were loaded into a 200keV Talos Arctica (Thermo Fisher) transmission electron microscope. 3,208 micrographs (Figure S1A) were acquired with a K2 Summit (Gatan) direct electron detector operating in super-resolution mode, using the Leginon automated data collection software^65^ at a nominal magnification of 36,000X (super-resolution pixel size of 0.575 Å/pixel; physical pixel size of 1.15 Å/pixel). Each micrograph was collected as dose-fractionated movie, where each movie comprised of 56 frames acquired over 14 s with a cumulative exposure of ∼58 electrons/Å^2^. A nominal defocus range of 0.6 *µ*m to 1.5 *µ*m was used for collecting the data.

### Image processing and 3D reconstruction

The super-resolution movie frames were first Fourier-binned 2 × 2 times to a pixel size of 1.15 Å/pixel, prior to dose-weighted frame alignment using MotionCor2^66^ implemented in the Appion^67^ image processing workflow. CTF parameters for the summed aligned micrographs were estimated using CTFFind4^68^ (Figure S1B) and only micrographs with confidence values above 90% were further processed. Particles were picked from these micrographs using the FindEM (Roseman et al., 2004) template-based particle picker in the Appion workflow, using selected 2D class averages from the previous Csy-Acr complex dataset as templates^24^. Coordinates from these picks were then imported into RELION 2.0^69^, and 1,543,677 particles were extracted with a box size of 288 pixels, which were binned by a factor of 2 (resulting box size 144 pixels, pixel size of 2.3 Å/pixel). These particles were then subjected to reference-free 2D classification (Figure S1C) within RELION 2.0, and a stack of 962,677 particles was obtained by selecting classes that represented different orientations and contained high-resolution features. These selected particles were subjected to 3D refinement (Figure S2A), using a 60 Å low passed filtered Csy-Acr map (EMD-8624) as an initial model. Particles from the 3D refinement were subjected to 3D classification without alignment and sorted into four classes. 743,861 particles belonging to two well-resolved 3D classes with the intact Cas8f C-terminal helix bundle were selected for further processing. Based on the × and y shifts associated with these particles, unbinned particles (box size 288 pixels, and pixel size of 1.15 Å/pixel) were extracted with re-centered coordinates. These particles were subjected to unmasked 3D refinement followed by another round of refinement with a soft edged 3D binary mask. The mask used for the refinement was generated using the volume from unmasked refinement run, that was expanded by 5 pixels with 8 pixels Gaussian fall-off smoothing. All subsequent masks that were used for downstream data processing were generated using the same procedure. The resulting reconstruction reported a resolution of 3.85 Å at a Fourier Shell Correlation (FSC) of 0.143. To further sort structural heterogeneity, particles from this 3D refinement were subjected to three class 3D classification without alignment. 291,227 particles from the best resolved 3D class of the full complex (containing the helix bundle of Cas8f) were further refined, resulting in a 3.4 Å resolution (at an FSC of 0.143) reconstruction (Figure S3G). Though the majority of this reconstruction presented well-defined structural details, the head, tail, and the helix bundle region of the Csy-DNA complex were poorly resolved due to intrinsic flexibility (Figure S2A and S2C).

In order to improve the quality of the map for the different regions of the Csy-DNA complex we used the signal-subtracted focused classification and refinement technique (Figure S2B) in RELION 2.1^24,70^. The whole complex was divided into three regions with some overlap between contiguous regions. These were the head-Cas8f helix bundle-Cas7f.1-Cas7f.2 subunits (region-1), the backbone comprising of all six Cas7f subunits and target DNA bound crRNA (region-2), and the tail-Cas7f.6 subunits (region-3). Each of the signal-subtracted particle stacks were subjected to independent 3D refinement and clustering (classification without alignment) runs, resulting in better quality map for each of the three regions. The final focused map for the head-Cas8f helix bundle-Cas7f.1 subunits, tail-Cas7f.6 subunits, and the backbone region were resolved to 3.3 Å, 3.2 Å and 3.1 Å (at 0.143 FSC value) (Figure S3F), respectively. In order to better facilitate model building of the full Csy-DNA complex, the three focused maps were aligned relative to each other, with the overlapping regions and the unsharpened non-focused reconstructed map of the full complex serving as guides and alignment references. A composite map was generated from the three focused maps by retaining the maximum valued voxel at each point, accomplished by using the “vop maximum” function in UCSF Chimera^71^ (Figure S2B). Local resolution estimations (Figure S1E) were calculated using the “blocres” function in the Bsoft suite^72^.

### Atomic model building

The atomic models for Cas5f, Cas8f, Cas6f and Cas7f from the Csy-Acr complex (PDB ID: 5UZ9) were used as initial template models for model building. These were individually rigid-body fitted into the reconstructed maps using the “fit map” function in UCSF Chimera^71^, and residue registers and backbone geometries were adjusted in Coot^73^. Models for the crRNA and DNA strands were also manually built into the map using Coot. This model underwent real-space refinement with rigid body fitting and simulated annealing in PHENIX^74^. The refined model was used as a seed for generating 200 models in Rosetta and the top scoring model was used for further refinement. Multiple rounds of refinement of the model was performed in PHENIX and Coot to fix the geometric and steric outliers, which were identified by MolProbity during validation. Once the major issues with the model were fixed, the final refinement iterations were carried out with secondary structure and Non-Crystallographic Symmetry (NCS) restrains. Regions of the map, particularly flexible loop regions and the R-loop of the target DNA could not be modeled due to lack of EM density. UCSF Chimera^71^ and ChimeraX^75^ were used for visualization and for generating all the figures for the maps and models (Figure S3A-E and Figure S3G).

### Electrophoretic Mobility Shift Assays (EMSA)

#### dsDNA binding assay

Binding assays were performed by incubating 0, 0.001, 0.01, 0.05, 0.1, 0.5, 1, 10, 100, 1000, 10,000 nM Csy complex with ʔ0.5 nM of 5’ ^32^P-labeled DNA oligonucleotides for 15 minutes at 37°C in reaction buffer (20 mM HEPES pH 7.5, 100 mM KCl, 5% glycerol, 1 mM TCEP). Reaction products were run on 6% polyacrylamide gels, which were dried and imaged with a phosphor storage screen (Kodak), then scanned with a Typhoon phosphorimager (GE Healthcare). Bands were quantified using ImageQuant software, and the percent DNA bound was plotted as a function of Csy complex concentration, then fit with a standard binding isotherm: Fraction DNA bound = [Csy complex]/(*K*D + [Csy complex])

#### Cas1-2/3 recruitment assay

5’ [^32^P]-labeled 80-base pair dsDNA (Table S1) was pre-incubated with 1 *µ*M Csy complex at 37°C in reaction buffer (20 mM HEPES pH 7.5, 100 mM KCl, 5% glycerol, 1 mM TCEP, 5 mM MgCl_2_, 75 [uni03BC]M NiSO_4_, 5 mM CaCl_2_, 1 mM ATP) for 15 minutes. Reactions were then moved to ice, and KCl concentration was increased to 300 mM to reduce non-specific interactions between DNA and Cas1-2/3. 1.85 nM, 5.5 nM, 16.6 nM, or 50 nM Cas1-2/3 was added to reactions, which were incubated for a further 5 minutes at 37°C. Reactions were separated by electrophoresis over native 4.5% polyacrylamide gels. Dried gels were imaged with a phosphor storage screen (Kodak), scanned with a Typhoon phosphorimager (GE Healthcare), and band intensities were quantified using ImageQuant software (GE Healthcare).

### Acknowledgments

We are grateful to Bill Anderson, The Scripps Research Institute (TSRI) Electron Microscopy facility manager, and Jean-Christophe Ducom at TSRI High Performance Computing for support during EM data collection and processing. G.C.L. is supported as a Searle Scholar, a Pew Scholar, by a young investigator award from Amgen, and the National Institutes of Health (DP2EB020402). Research in the Wiedenheft lab is supported by the National Institutes of Health (P20GM103500, P30GM110732, R01GM110270, R01GM108888 and R21 AI130670), the National Science Foundation EPSCoR (EPS-110134), the M. J. Murdock Charitable Trust, a young investigator award from Amgen, and the Montana State University Agricultural Experimental Station (USDA NIFA). Computational analyses of EM data were performed using shared instrumentation at TSRI funded by NIH S10OD021634.

### Data and materials availability

All the cryo-EM maps were deposited in the Electron Microscopy Data Bank under accession code EMD-9191. The associated atomic coordinates were deposited into the Protein Data Bank with accession code 6MPU.

## Supplementary Materials

**Figure S1.**
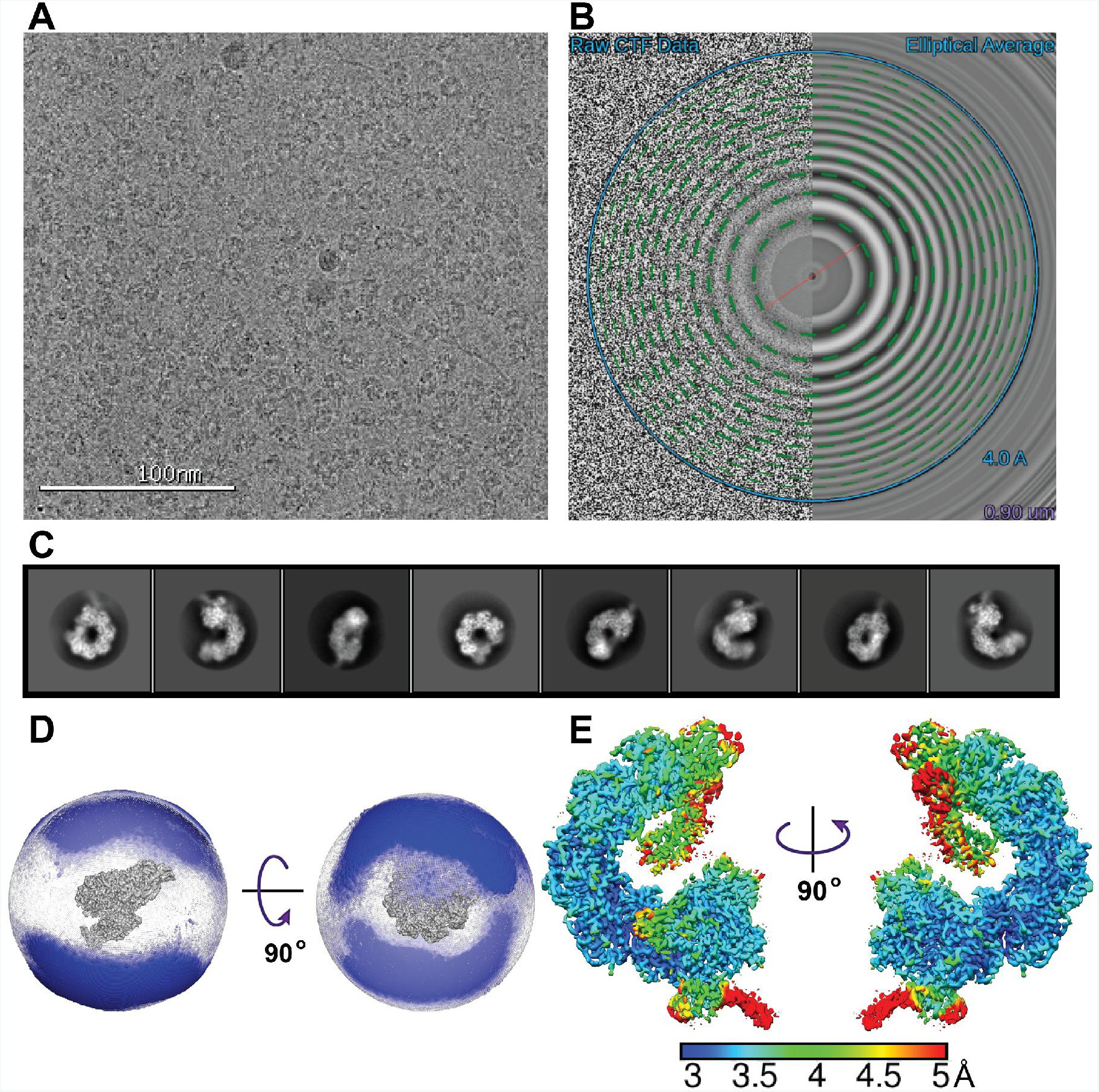
Cryo Electron Microscopy of the target DNA bound Csy complex. (A) A representative cryo-EM micrograph of the Csy-DNA complex in vitreous ice. The ring-shaped features are the individual particles of the complex. (B) Fourier transform of the micrograph shown in (A), with the Contrast Transfer Function (CTF) estimation. The Thon Rings can be fitted with high confidence (green dotted circles) beyond 4 Å resolution (blue circle). (C) Representative reference-free 2D class averages of the Csy-DNA complex showing different orientations of the complex in ice and also high-resolution features. (D) Euler angle distribution of the particles that contributed to the final 3D reconstruction of the complex (map shown in gray at the center of the spherical plots). The angular assignments are denoted by the position of the blue spheres relative to the map in the center, and the radius of the spheres correspond to the number of particles in that angular orientation. (E) The final 3D map colored based on local resolution estimation using “blocres” function of the Bsoft suite^72^. The majority of the map has been resolved to between 3-4 Å resolution, with peripheral regions of the flexible head, tail and DNA duplex not as well resolved.

**Figure S2.**
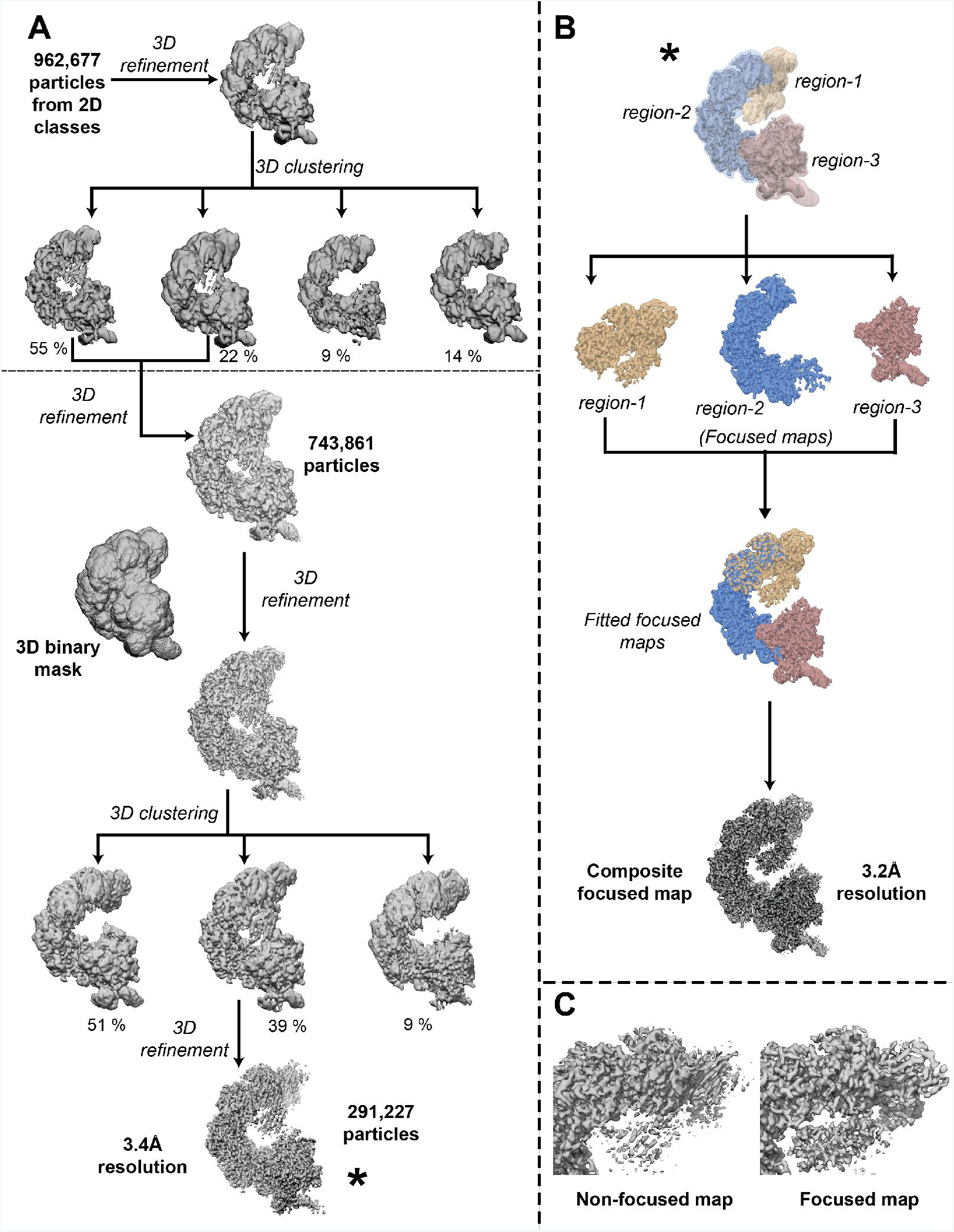
Schematic of the 3D reconstruction and data processing workflow. (A) Initial steps of cryo-EM data processing were carried out with binned pixel size of 2.3 Å/pixel and box size of 144 pixels. Particles selected from well-resolved 2D classes were subjected to initial 3D processing. Subsequent processing was carried out with unbinned particles with box size of 144 pixels, pixel size of 1.15 Å/pixel. The thin horizontal dotted line separates processing steps using binned and unbinned data. All 3D classification steps were carried out without alignments and are referred to as “3D clustering”. The percentage value below 3D classes are the number of particles from the preceding stack that contributed to the class. The final 3D reconstruction and the particle stack (indicated with *) were subjected to signal subtracted focused processing to improve the resolutions of the different sub-regions of the complex. (B) The Csy-DNA complex was divided into three contiguous regions (detailed description in Methods) (colored yellow, blue and brown). Each region was subjected to focused analysis and the resulting reconstructions are shown. These reconstructions were then fitted relative to each other and a composite stitched map was created using the “vop maximum” function in UCSF Chimera^71^ (C) Structural features were better resolved in the composite focused map as compared to the non-focused map. As an example, comparison between the non-focused (right side) and focused (left side) maps for “region-1” (head-Cas8f helix-bundle-Cas7.1f-Cas7.2f subunits) are shown.

**Figure S3.**
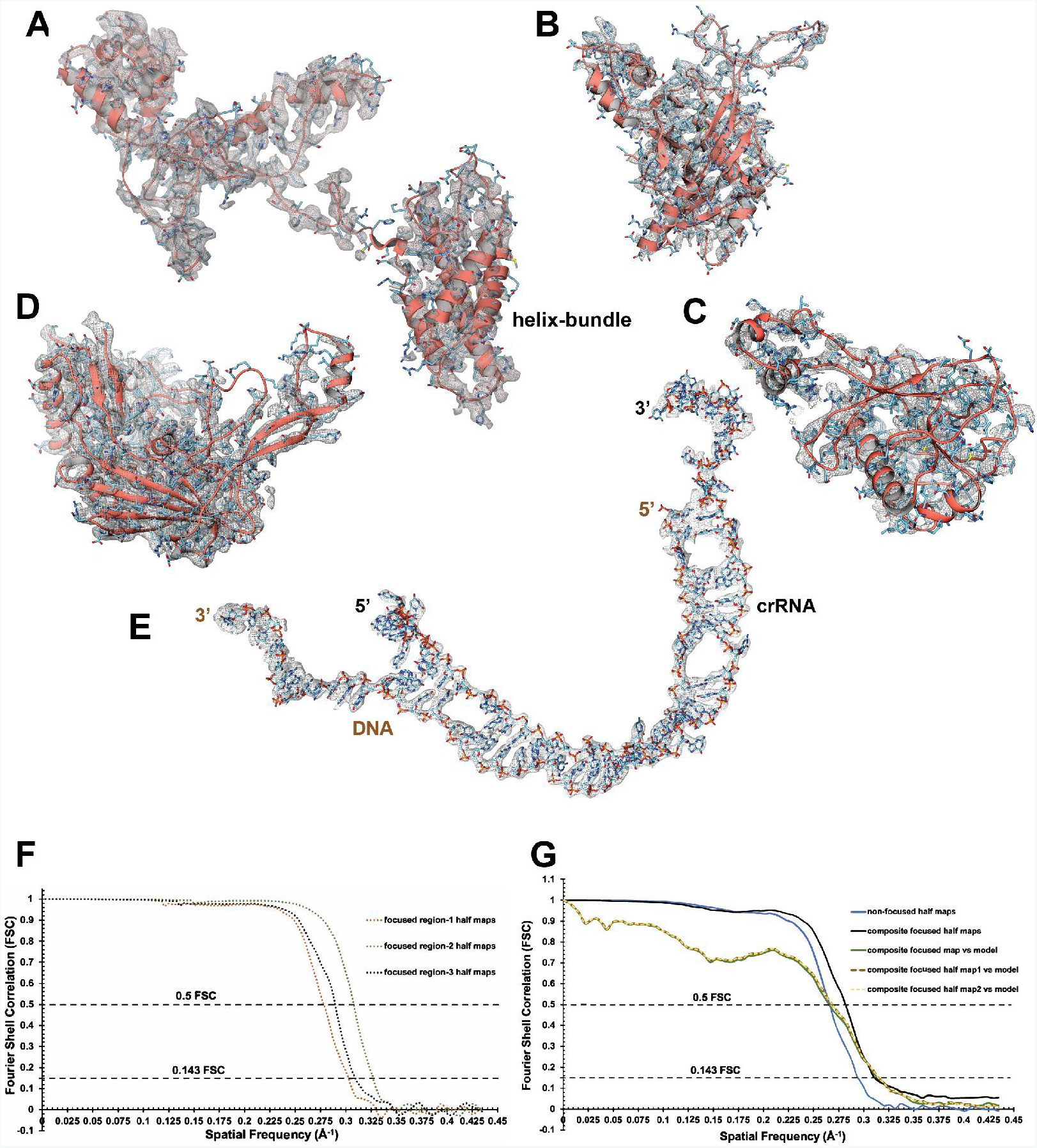
Cryo-EM density and atomic models of subunits. EM density of the subunits are shown as gray mesh with the built-in atomic models. (A) Cas 8f subunit map and model with the C-terminal helix bundle. (B) Cas5f subunit map and model. (C) Cas6f subunit map and model. (D) One of the representative Cas7f subunit map and model. (E) Density map and atomic model of the crRNA hybridized with the complimentary strand of the target DNA. (F) Fourier Shell Correlation (FSC) curves between focused “region-1” half maps (orange dotted), focused “region-2” half maps (green dotted), and focused “region-3” half maps (black dotted). Detailed description of the three regions are provided in methods. (G) Fourier Shell Correlation (FSC) curves between reconstructed non-focused half maps (blue), focused composite half maps (black), summed composite map vs model (green), and model vs one half focused composite map (brown dashed), and another half focused composite map (yellow dashed).

**Figure S4.**
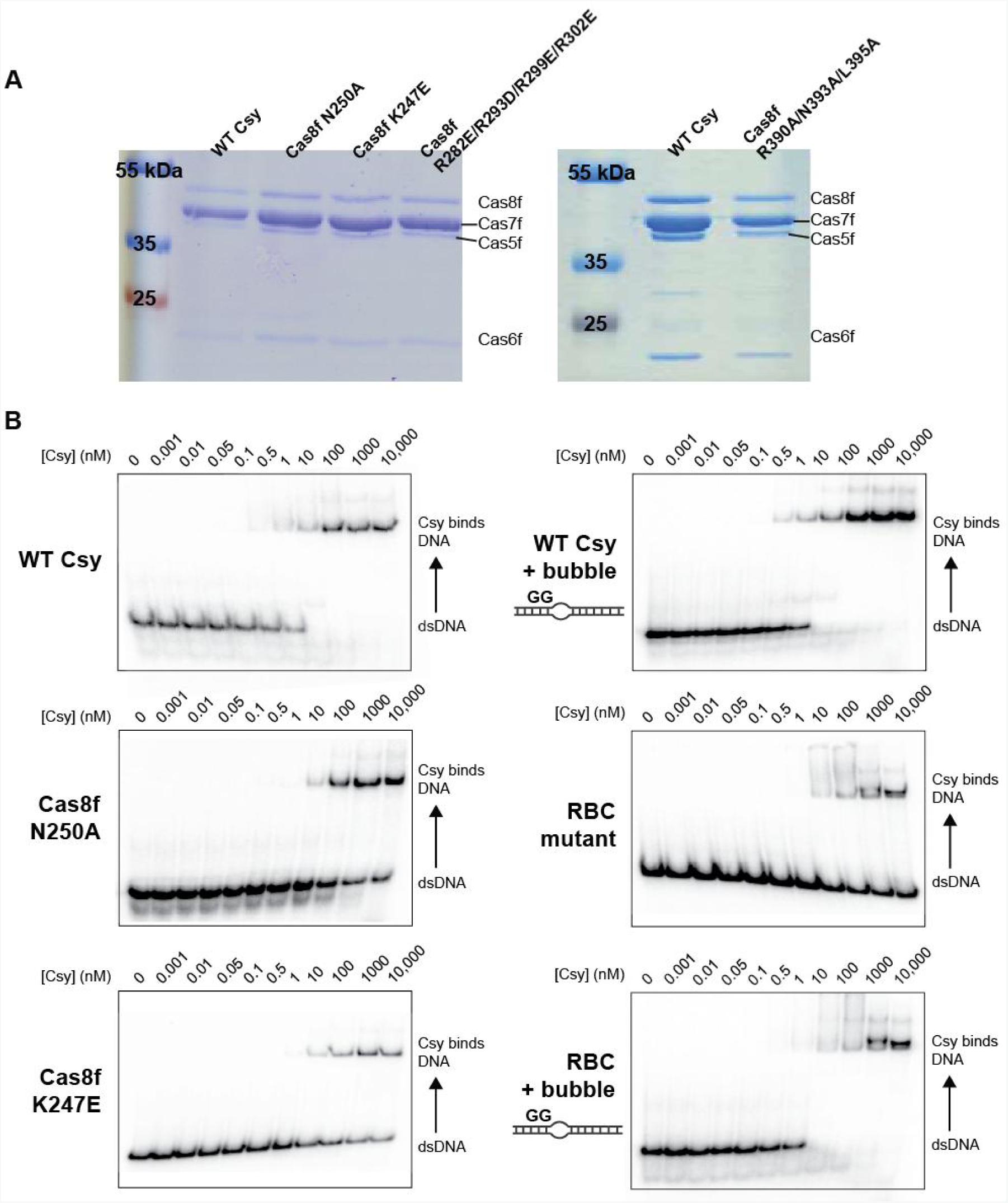
Purification of and activity assays for mutant Csy complexes. (A) Coomassie blue-stained SDS-PAGE gels including WT and mutant Csy complexes. (B) DNA binding assays were performed by incubating a concentration gradient (0, 0.001, 0.01, 0.05, 0.1, 0.5, 1, 10, 100, 1453720, 10,453720 nM) of Csy complex with ¡0.5 nM of 5’ ^32^P-labeled dsDNA. Gels are representative of replicate experiments.

**Figure S5.**
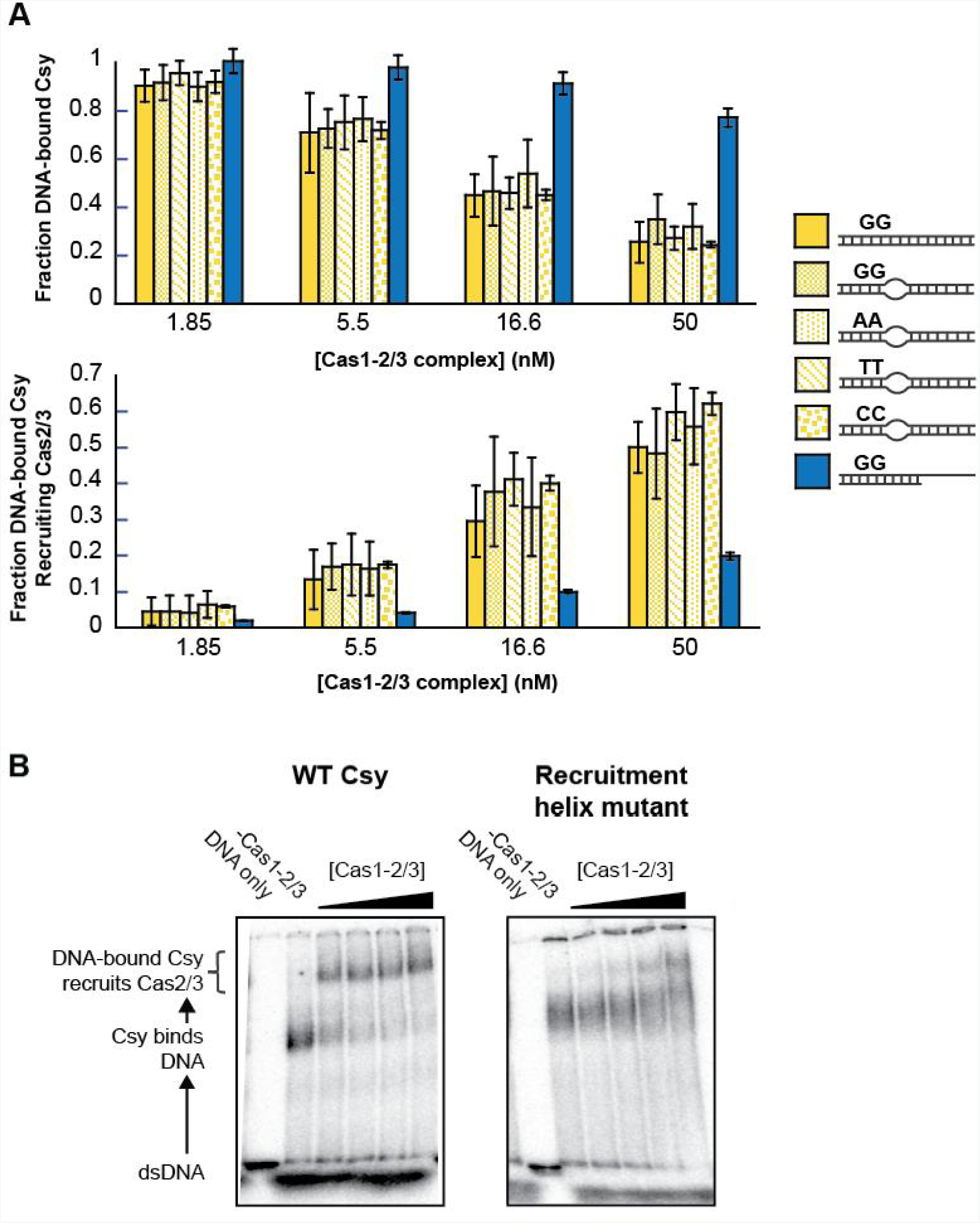
Cas2/3 recruitment requires a complete R-loop and the “recruitment helix” on Cas8f. Electrophoretic mobility shift assays (EMSAs) were performed with radiolabeled dsDNA substrates, purified Csy complex and increasing concentrations (1.85 nM, 5.5 nM, 16.6 nM or 50 nM) of the Cas1-2/3 complex. (A) Quantification of EMSAs show a Cas1-2/3-dependent decrease in dsDNA-bound Csy complex, and corresponding increase in dsDNA-Csy-Cas2/3 supercomplex. This was seen for all DNA substrates tested except the partial duplex. Representative gels shown in Figure 4C. (B) Mutations in the “recruitment helix” of Cas8f impair Cas2/3 recruitment.

**Table S1.**
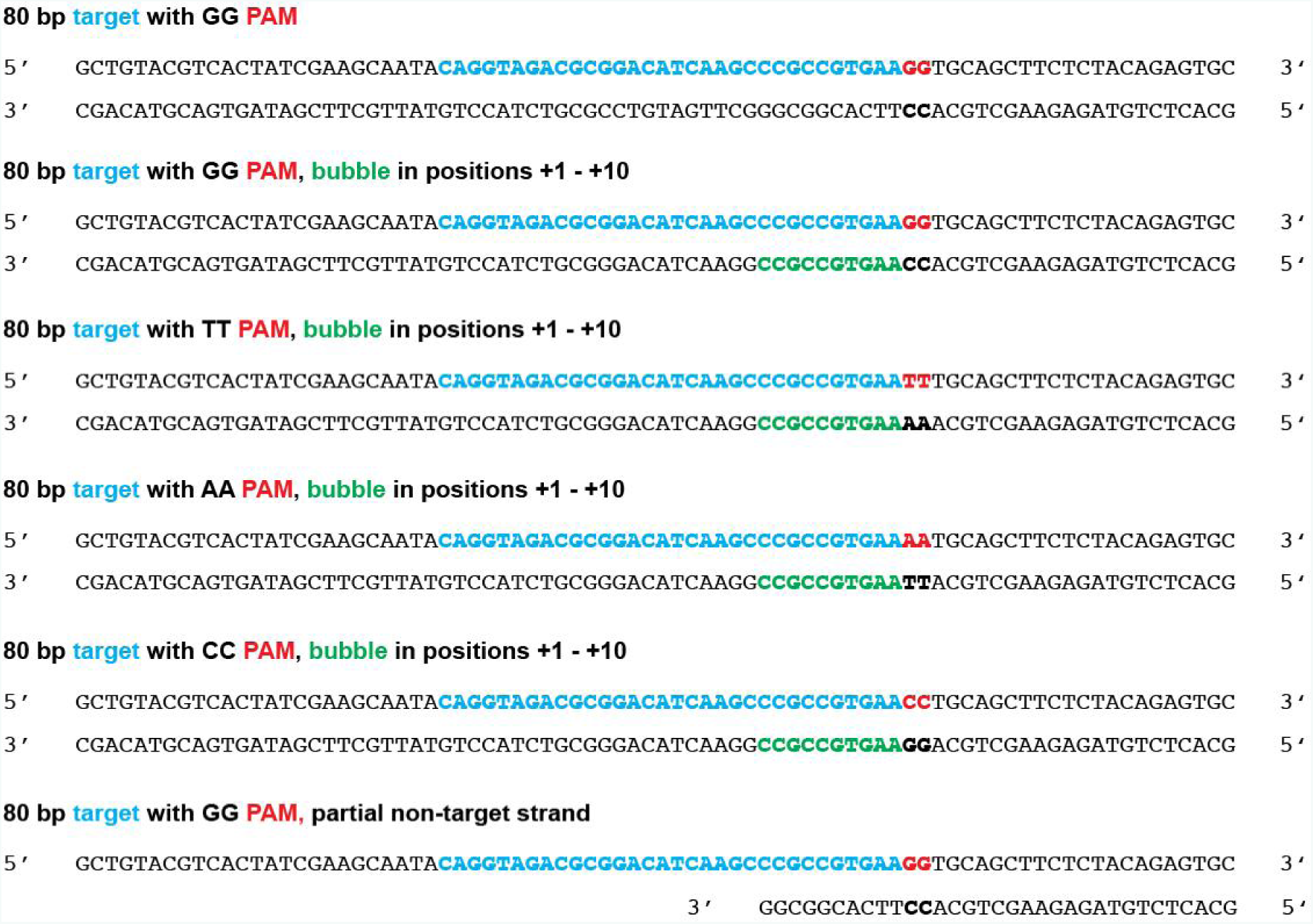
DNA oligonucleotides

**Table S2.**
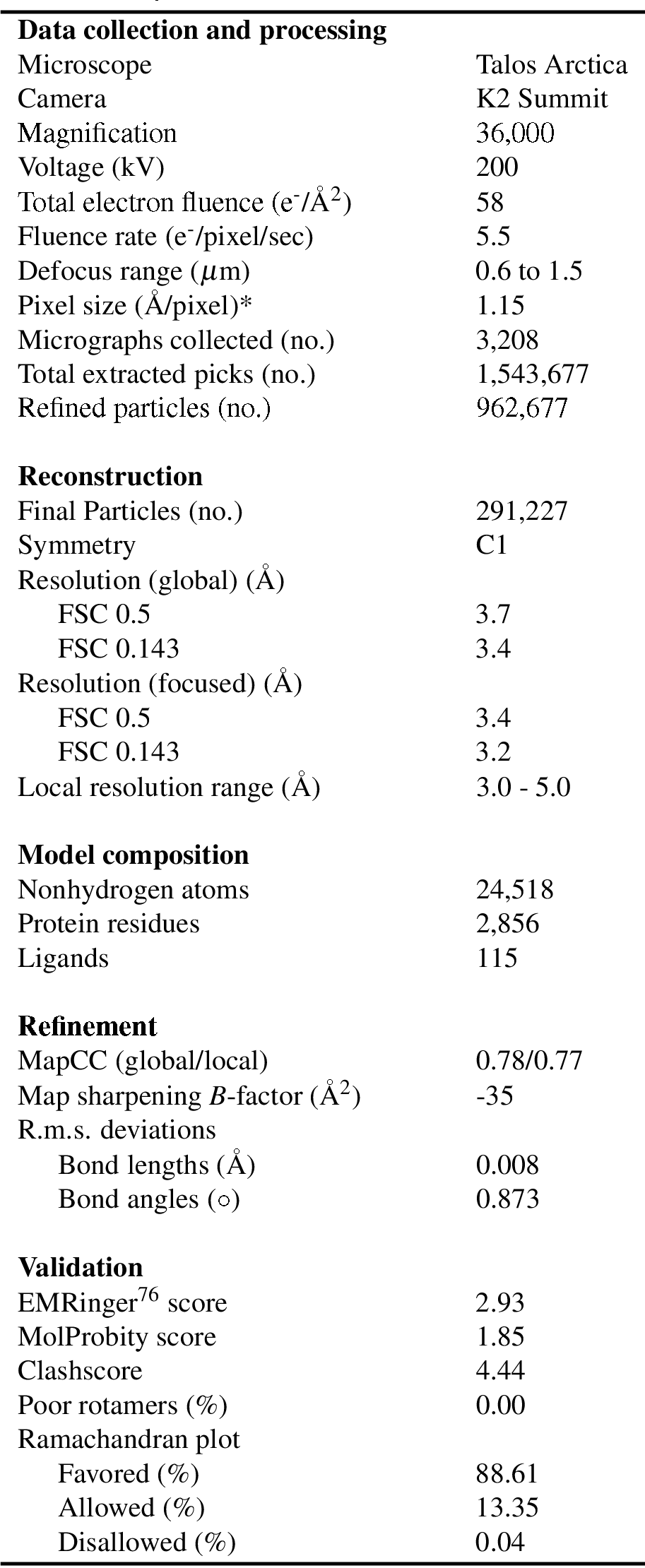
Cryo-EM data collection, refinement, and validation statistics

* Calibrated pixel size at the detector

Movie S1. Atomic model of the Csy complex binding dsDNA and structural rearrangements. Atomic models of unbound Csy complex (PDB ID: 6B45), Csy bound to a partially duplexed dsDNA (PDB ID: 6B44), and Csy bound to a complete dsDNA target (PDB ID: 6MPU) were used to generate a linear interpolation representing the conformational changes that the Csy complex undergoes during DNA binding. The Csy complex initially engages DNA through non-sequence-specific electrostatic interactions, followed by sequence-specific interactions with the protospacer adjacent motif (PAM). Incoming DNA is immobilized in a vice formed by the N-terminal domain of the large subunit (Cas8f) and the opposing face of the terminal backbone subunit (Cas7f.6).

Movie S2. Details of the R-loop interactions with the Csy complex. Residues in Cas8f detect the PAM in the minor groove, locally distorting the DNA duplex and facilitating strand invasion. The complementary DNA strand hybridizes with the crRNA guide, forming an R-loop that is stabilized by positively-charged residues in an “R-loop binding channel” that terminates near the 3’ end of the crRNA spacer. Formation of the complete R-loop is critical for rotation of the C-terminal helical bundle of Cas8f and recruitment of the trans-acting nuclease-helicase Cas2/3.

Movie S3. Structural similarity between AcrIF3 and the Cas8f helical bundle. A virally-encoded anti-CRISPR protein (AcrIF3) is a molecular mimic of the Cas8f helical bundle, and comparison of the two structures reveals the 180-degree rotation of the helical bundle of Cas8f exposes a “nuclease recruitment helix.” Collectively, the model explains how the Csy complex coordinates nuclease recruitment to *bona fide* dsDNA targets.

